# The brain selectively tunes to unfamiliar voices during sleep

**DOI:** 10.1101/2021.08.26.457494

**Authors:** Mohamed S. Ameen, Dominik PJ Heib, Christine Blume, Manuel Schabus

**Author notes:** These authors contributed equally (Dominik PJ Heib, Christine Blume).

## Abstract

The brain continues to respond selectively to environmental stimuli even during sleep. However, the functional role of such responses, and whether they reflect information processing or rather sensory inhibition is not fully understood.

Here, we presented 17 human sleepers (14 females) with their own name and two unfamiliar first names, spoken by either a familiar voice (FV) or an unfamiliar voice (UFV), while recording polysomnography during a full night’s sleep. We detected K-complexes, sleep spindles, and micro-arousals, and then assessed event-related potentials, oscillatory power as well as intertrial phase synchronization in response to the different stimuli presented during non-rapid eye movement (NREM) sleep.

We show that UFVs evoke more K-complexes and micro-arousals than FVs. When both stimuli evoke a K-complex, we observed larger evoked potentials, higher oscillatory power in the high beta (>16Hz) frequency range, and stronger time-locking in the delta band (1-4 Hz) in response to UFVs relative to FVs. Crucially, these differences in brain responses disappear when no K-complexes are evoked by the auditory stimuli.

Our findings highlight discrepancies in brain responses to auditory stimuli based on their relevance to the sleeper and propose a key role for K-complexes in the modulation of sensory processing during sleep. We argue that such content-specific, dynamic reactivity to external sensory information enables the brain to enter a ‘sentinel processing mode’ in which it engages in the many important processes that are ongoing during sleep while still maintaining the ability to process vital information in the surrounding.

**Significance statement:** Previous research has shown that sensory processing continues during sleep. Here, we studied the capacity of the sleeping brain to extract and process relevant sensory information. We presented sleepers with their own names and unfamiliar names spoken by either a familiar (FV) or an unfamiliar voice (UFV). During non-rapid eye movement (NREM) sleep, UFVs elicited more K-complexes and micro-arousals than FVs. By contrasting stimuli which evoked K-complexes, we demonstrate that UFVs triggered larger evoked potentials, stronger time-locking in the delta (1-4Hz) band, and higher oscillatory power (>16Hz) relative to FVs. These differences in brain responses disappeared when no K-complexes were evoked. Our results suggest a pivotal role for K-complexes in the selective processing of relevant information during NREM sleep.

## 1. Introduction

During sleep, the brain continues to respond to auditory stimuli in a selective fashion (Portas et al., 2000; Andrillon et al., 2016; Blume et al., 2017; 2018). Previous studies have demonstrated that, for instance, the subject’s own name (SON) evokes stronger brain responses than other names during sleep (Oswald et al., 1960; Perrin et al., 1999; Pratt et al., 1999). Recently, Blume et al. (2018) showed that during all stages of sleep, brain responses to SON and other unfamiliar names (UNs) did not differ; however, names uttered by an unfamiliar voice (UFV) evoked stronger brain responses as compared to a familiar voice (FV).

The discrepancy in brain responses to different stimuli implies the presence of an initial, presumably low-level, sensory processing during sleep that enables the brain to differentiate between sensory signals (Blume et al., 2018). However, knowledge about the functions of such responses is still lacking. That is, the selective brain responses to specific sounds during sleep might reflect inhibitory processes that protect sleep from disruptions. Conversely, they might indicate further, higher level, processing that ensures the connectedness of the sleeping brain to the surrounding.

In this study, we investigated the purpose of such selective brain responses to sounds presented during non-rapid eye movement (NREM) sleep. We focused on sleep-specific events that have been previously linked to information processing, sensory inhibition, or both. We focused on three cardinal sleep-specific events, namely, the K-complex (KC), sleep spindles, and micro-arousals.

KCs are ~1Hz oscillations and a hallmark of N2 sleep (Loomis et al., 1938, Colrain, 2005; Halász, 2005). KCs either occur spontaneously or evoked by sensory stimuli. Spontaneous KCs appear in the electroencephalography (EEG) signal as well-defined sharp negative wave followed by a positive component with a total duration of at least 0.5s (Rechtschaffen and Kales, 1968; Hori et al., 2001). Following sensory perturbation, KCs appear to have two main components; a negative deflection at ~550ms (N550) followed by a longer-lasting positive wave at ~900ms (P900) (Bastien & Campbell, 1992, Cote et al., 1999; Colrain, 2005; Halász, 2005). Some studies have considered an early positive peak that appears around 200ms (P200, Laurino et al., 2014; 2019) and another negative peak around 350ms (N350, Bastien & Campbell, 1992, Cote et al., 1999) to be parts of the KC, albeit these potential can occur without a KC being elicited. Relevant stimuli have higher propensity to trigger KCs (Halász, 2005). Recent theories suggest that KCs can serve both sleep-protecting as well as arousal-inducing processes (Halász, 2005; Jahnke et al., 2012; Forget et al., 2014; Laurino et al., 2014; Blume et al., 2017; 2018; Legendre et al., 2019; Latreille et al., 2020).

Sleep spindles are also characteristic of N2 sleep. Spindles are thalamocortical oscillations of 11-15 Hz that last around 0.5 to 2s (De Gennaro and Ferrara, 2003; Fernandez and Lüthi, 2020) that can be triggered by sensory stimuli (Antony and Paller, 2017). They have been repeatedly shown to inhibit sensory processing during sleep (McCormick & Bal, 1994, Schabus et al., 2012; Blume et al., 2018; Fernandez and Lüthi, 2020). However, recent work challenges this notion (Sela et al., 2016) and even associates spindles with the processing of memory-related sounds presented during NREM sleep (Cairney et al., 2018).

Finally, micro-arousals are abrupt shifts in the EEG signal towards theta, alpha or higher theta, alpha and/or high beta (>16 Hz) frequencies (Halász et al., 1979; American Sleep Disorders Association, 1992; Halász et al., 2004) that appear in all sleep stages and are considered windows of information processing during sleep (dos Santos Lima et al., 2019; Halász et al., 2004; Halász, 2005). Micro-arousals are usually preceded by KCs (Colrain 2005; Halász, 2005), yet they have been shown to be correlated with a lower incidence of sleep spindles in the preceding 10s of EEG signal (Ehrhart et al., 1981).

Here, we re-analysed the dataset used in Blume et al. (2018) who recorded polysomnography while presenting SONs and two UNs spoken by either a FV or an UFV during a whole night of sleep. We detected KCs, spindles, and micro-arousals in response to these sounds during NREM sleep and hypothesized that the selective auditory-evoked responses during NREM sleep support the extraction and processing of relevant sensory information.

## 2. Materials and methods

### 2.1. Participants

We recruited 20 healthy participants with no reported history of neurological or psychological problems as well as no reported sleep disorders. However, one participant dropped out after the adaptation night and we had to exclude two participants due to technical problems during EEG acquisition. Therefore, we performed the analyses we report here on 17 participants (14 females) with a median age of 22.6 years (SD = 2.3). Before beginning the experiment, participants signed written informed consents. The experiment was approved by the ethics committee of the University of Salzburg.

### 2.2. Experimental design

Before the start of the experiment, participants were advised to maintain a regular sleep/wake cycle (~8h of sleep) for at least four days which we monitored via actigraphy (Figure 1A). Subsequently, participants spent two nights in the sleep laboratory of the University of Salzburg. The first night was an adaptation night, during which we recorded polysomnography (PSG) data with no auditory stimulation. The second night was an experimental night, during which we recorded PSG data while presenting sounds via loudspeakers throughout the night. In both nights, participants were tested during wakefulness before and after sleep. Briefly, the wakefulness testing consisted of two sessions, a passive-listening session and an active-listening session. Passive listening entails that the participants had to listen to the repeatedly presented auditory stimuli; while active listening means that, they had to count the number of presentations of one specific stimulus chosen by the experimenters. Before the wakefulness testing, participants were stimulated with either a bright (blue-enriched) light or an inactive (sham) light for one hour. The order of the light-stimulation conditions was counterbalanced between the adaptation and the experimental nights across participants. However, the light condition is irrelevant to this study, as we grouped our data over both light conditions. For the purpose of this paper, we will focus primarily on the sleep part of the experimental night. For more details on the wakefulness part of the experiment, please refer to Blume et al. (2018). During the experimental night, participants went to bed around their habitual bedtime (8:30pm – 11:30pm). Time in bed was approximately 8 hours. After 8 hours, we waited for light NREM or REM sleep before waking up the participants (median sleep duration = 480 ± 2.5 minutes). The auditory stimulation started directly after participants went to bed and continued throughout the whole night. We presented auditory stimuli continuously for 90 minutes (*Stimulation periods*) then paused the presentation for 30 minutes (*No-stimulation periods*) to allow for periods of undisturbed sleep. This resulted in a 120-minute cycle that we repeated four times throughout the night (Figure 1B).

**Figure 1.**
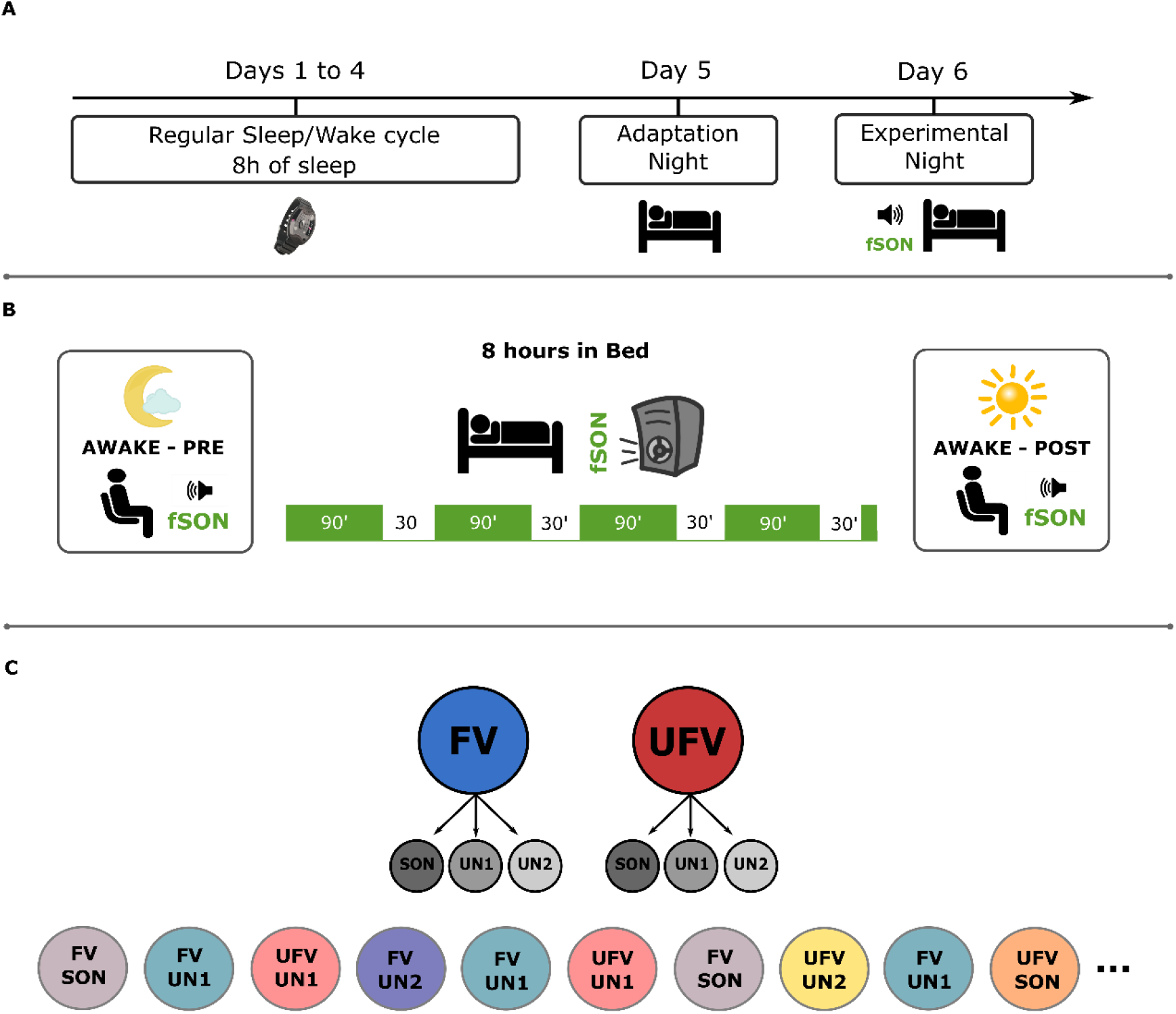
Experimental design. A) Protocol: participants were invited for an initial screening interview during which they were given the wrist-worn actigraphy and advised to keep a regular sleep-wake cycle. Participants slept in the sleep laboratory during two nights: an adaptation night during which PSG was recorded but no stimuli were presented, and an experimental night during which we recorded PSG and presented auditory stimuli throughout the night. B) Procedure of the testing night: participants slept for approximately eight hours with PSG and auditory stimulation. All participants went through a stimulation session in the evening before sleep; however, this session is irrelevant to the current manuscript. The auditory stimuli started directly after going to bed and continued for 90 minutes, then paused for ~30 minutes to allow for a period of undisturbed sleep. This cycle was repeated four times accounting for the whole duration of the night. C) (Top) Stimuli: We presented the subject’s own name (SON) and two unfamiliar names (UN1 & UN2) spoken by either a familiar voice (FV) or an unfamiliar voice (UFV). (Bottom): an exemplary sequence of stimulus presentation. Stimuli were presented in a pseudo-random order; each stimulus was presented 690 times and the inter-stimulus intervals ranged between 2800ms to 7800ms.

### 2.3. Stimuli

We presented six different auditory stimuli (Figure 1C) that we personalized for each participant. The stimuli were the subject’s own name (SON) and two unfamiliar names (thereafter referred to as UNs) spoken by either a familiar voice (FV) or an unfamiliar voice (UFV). A FV was the voice of someone close to the participant, for example one of the parents. An UFV was the voice of someone unknown to the participant. We did not control for the sex of the voices, but they were matched, that is, the familiar and unfamiliar voices were always both either male or female. We chose UNs that matched the SON in the number of syllables and the frequency of occurrence in the population. The volume for stimulus presentation was adjusted individually for each participant so that the participant could clearly hear the stimulus and still be able to fall asleep. Each stimulus was presented 690 times and the mean duration was 808ms ± 110ms. Stimuli were presented in a pseudo-random order and no stimulus was presented twice in a row. The inter-stimulus intervals during sleep were jittered between 2800ms to 7800ms in 500ms steps. Stimulus pre-processing, i.e. denoising and normalization, was done using Audacity^©^ software (http://audacityteam.org/). Stimulus delivery was controlled by Matlab^®^ (Mathworks, Natick, USA).

### 2.4. Brain Data Acquisition

We recorded ongoing brain activity using a high-density EEG 256-channel GSN HydroCel Geodesic Sensor Net (Electrical 478 Geodesics Inc., Eugene, Oregon, USA) and a Net Amps 400 amplifier. The PSG recordings included two electrooculography (EOG) and two chin electromyography (EMG) channels. Data was acquired at a sampling rate of 250 Hz and Cz served as the online reference.

### 2.5 Sleep staging

Sleep staging was performed on 30s epochs using the computer-assisted sleep classification system developed by the SIESTA group (Somnolyzer 24×7; The SIESTA Group Schlafanalyse GmbH, Vienna, Austria; Anderer et al., 2005; Anderer et al., 2010) following the standard criteria recommended by the American Association for Sleep Medicine (AASM, American Academy of Sleep Medicine & Iber, 2007). We have previously shown that level of agreement between this algorithm and expert human scorers is similar to the level of agreement between human experts (Ameen et al., 2019).

### 2.6. The detection of sleep microstructures

#### 2.6.1. KC detection

We detected KCs automatically with a wavelet-detection algorithm that was developed by the SIESTA group (The SIESTA Group Schlafanalyse GmbH., Vienna, Austria). The development and validation procedures have been described in detail in Parapatics et al. (2015) and Schwarz et al. (2017), respectively. Briefly, 12 experienced human scorers visually scored KCs in 873 10-minute-epochs of PSG recordings from 189 control subjects and 90 patients. The features of the visually scored KCs were used to set the criteria for detection as well as to create a template KC that works as a gold standard for the automatic detection. The detection itself is a two-step process; first, the algorithm detects possible KCs via an approach that combines a matched-filtering detection method and a slow-wave detection-based method (Woertz et al., 2004). The detection criteria for possible KCs are: 1) a minimum negative-to-positive peak-to-peak amplitude of 50 μv, and 2) a duration between 480ms and 1500ms. Second, all possible KCs are matched to the prototypical KC template via wavelet analysis and the results are submitted to a linear discriminant analysis (LDA) to select only “real” KCs. For our analysis, we considered real KCs to be events that have an LDA score (how likely a specific EEG segment is a KC) of 0.8 or higher. This LDA value corresponds to 61.87% ± 9.14 of all detected KCs and a mean correlation to the template of 0.87 ± 0.007 over all subjects. Before running the detection algorithm, raw data were down-sampled to 128Hz and re-referenced to the contralateral mastoid. The algorithm detects KCs at C3 and C4. We only report results from C3 as the detections were similar between C3 and C4. We only considered events that occurred during N2 and N3 sleep. For N3 detections, however, we applied a more strict amplitude criterion as we only selected events with a peak-to-peak amplitude higher than 75μv (Cote et al., 1999; Nir et al., 2011). Fig. 2 demonstrates the LDA distribution of the detected KCs as well as some examples of the detected events. We defined evoked KCs as those events which occurred in the 2000ms post-stimulus-onset window.

**Figure 2.**
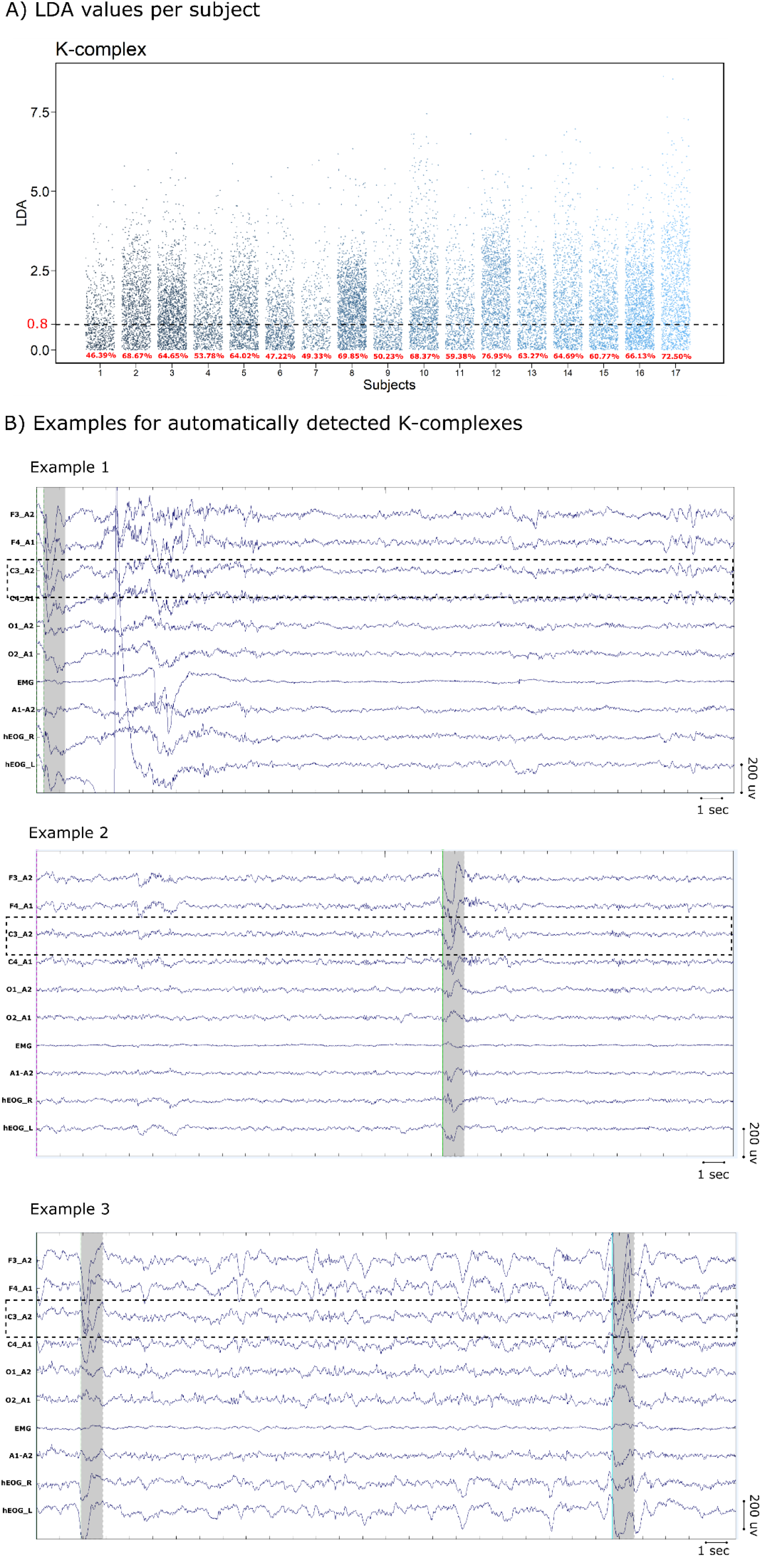
K-complex detection and examples. A) Linear discriminant analysis (LDA) values of the all detected events for all subjects. The dashed line represent our minimum cut-off at LDA value of 0.8. The percentage of the events used in our analyses from all the detected events for each subject are indicated in red at the bottom of the plot. B) Examples of the detected K-complexes. We show 30s epochs and the standard EEG montage that is used for sleep staging as well as event detections. Specifically, we used channels (from top to bottom): F3, F4, C3, C4, O1, and O2 all referenced to the contralateral mastoid. Moreover, we show one EMG and two EOG channels as well as the average of both mastoids. Examples 1 and 2 are events detected in N2 sleep from one subject and example 3 is a N3 epoch from a different subject.

#### 2.6.2. Spindle detection

Sleep spindles were detected using an algorithm developed by the SIESTA group (ASK analyser, The Siesta Group Schlafanalyse GmbH., Vienna, Austria, Gruber et al., 2015). First, we filtered the raw data between 11 and 16 Hz and then detected spindle events at frontal (F3, F4, Fz) and central leads (C3, C4 & Cz) channels re-referenced to the average of mastoids. We used the criteria described in Schimicek et al. (1994). Only events with an amplitude >12μv and duration between 500ms and 2000ms were considered. Further validation of the detected spindles was done using LDA in which the detected spindles were compared to a template that is generated based on the visual scoring of sleep spindles in 8730 minutes of PSG data from 189 healthy participants and 90 participants with sleep disorders. For our analyses, we only considered events that occurred during N2 and N3 sleep with an LDA score of 1.7 or higher (for more details see: Anderer et al., 2005). In order to identify the frequency of each spindle event, the algorithm preforms period-amplitude analysis of the band-pass filtered signal in the time domain. We subdivided spindles into slow (11-13 Hz) and fast (13-15Hz) spindles based on the dichotomy in their topography and functions (Schabus et al., 2007). We report results from fast spindles detected at C3 and slow spindles detected at F3.

#### 2.6.3. Micro-arousal detection

We detected micro-arousals semi-automatically using an algorithm developed by the SIESTA group (The Siesta Group Schlafanalyse GmbH., Vienna, Austria), and have been described in details in (Anderer et al., 2010). Briefly, the algorithm was developed using the scoring of 12 PSG recordings by six independent experts. It incorporates information from central and occipital channels. First, the algorithm compares the absolute and relative power of nine frequency bands including theta, alpha and high beta (> 16 Hz) frequencies between a 3-second test window and a moving ten-second baseline via a series of LDA separately for each channel. Second, the start and end of the micro-arousal events are determined by combining the posterior probabilities of all channels so that the number of micro-arousals per total sleep time is the same for both automatic and visual detections. The algorithm detects micro-arousals in all sleep stages, however, for the purpose of this study; we selected micro-arousals that occurred during N2 and N3 only. For examples of the detected micro-arousals see: Fig. 3).

**Figure 3.**
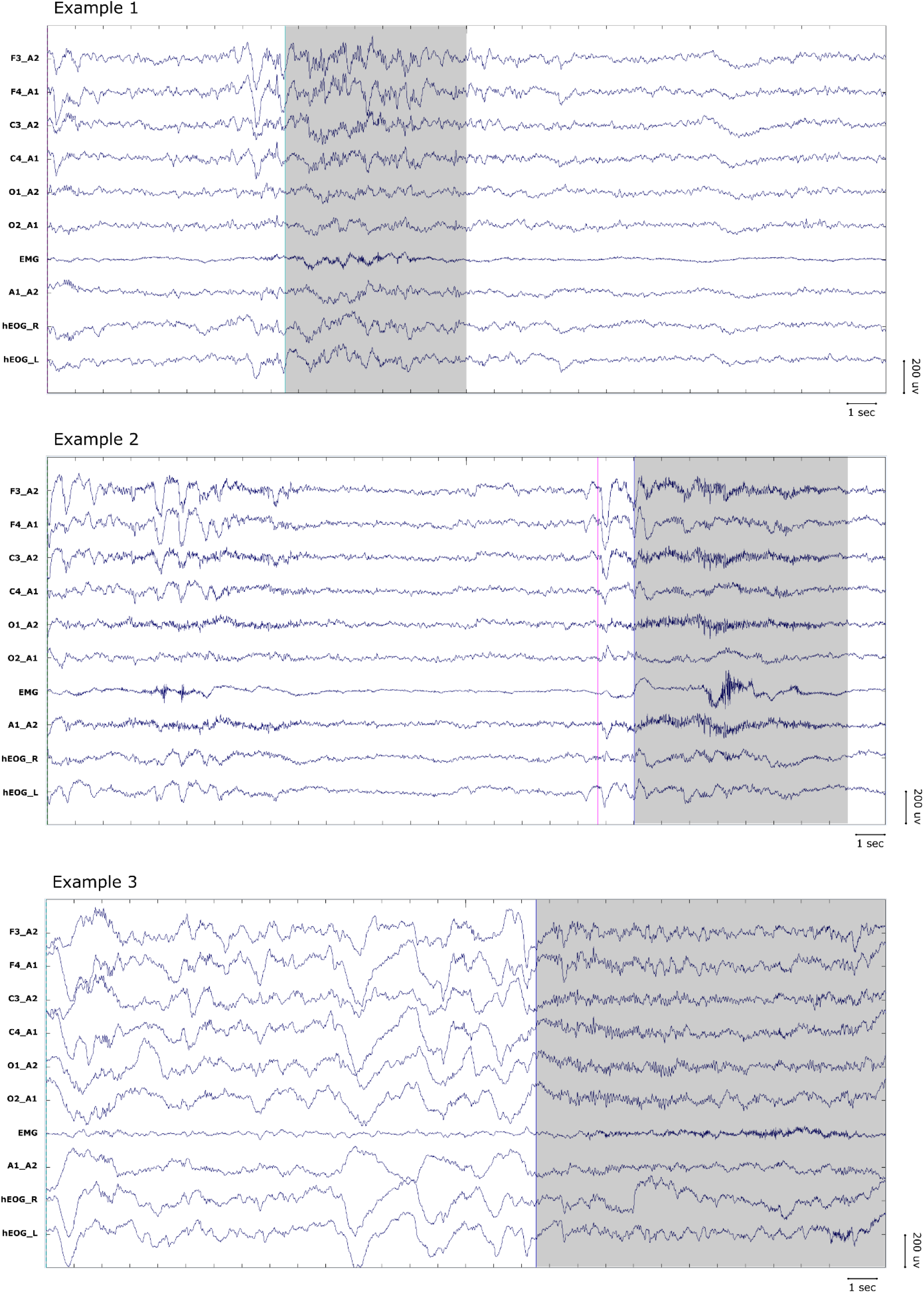
Examples of the detected micro-arousals. We show 30s epochs and the standard EEG montage that is used for sleep staging as well as event detections. Specifically, we used channels (from top to bottom): F3, F4, C3, C4, O1, and O2 all referenced to the contralateral mastoids. Moreover, we show one EMG and two EOG channels as well as the average of both mastoids. Examples 1 and 2 are events detected in N2 sleep from one subject, example 2 shows a micro-arousals that is precede by a K-complex. Example 3 is a N3 epoch from a different subject.

#### 2.6.4. The detection of transient microstates (micro-measures of alertness)

The detection of sleep microstates provides a more fine-grained scoring that detects transient changes in sleep architecture in 4s epochs rather than 30s, using an algorithm described in Jagannathan et al. (2018). The algorithm uses the Hori scale (Tanaka et al., 1996) to classify the epochs into either awake, drowsy (N1), or asleep (N2) based on the signal from a subset of 14 electrodes distributed over frontal, central, parietal, occipital and temporal regions. We extracted stimuli from N2 sleep only and removed stimuli with inter-stimulus intervals of less than 4000ms. Then, we filtered the data between 0.1 and 30Hz before running the algorithm on an equal number of epochs in all conditions.

### 2.7. EEG pre-processing and analyses

#### 2.7.1. Pre-processing

We performed all the pre-processing steps in EEGLAB v14.1.1b (Delorme & Makeig, 2004). First, we excluded face and neck channels and down-sampled the raw data from 183 EEG channels to 128 Hz. Then, we filtered the data between 0.1 and 40 Hz using a Butterworth bandpass filter. We performed bad channels rejection and interpolation as well as re-referencing to an average reference using the PREP pipeline described in Bigdely-Shamlo et al. (2015). Finally, we performed independent component analysis using the Adaptive Mixture Independent Component analysis (AMICA) toolbox and visually detected and discarded eye and muscle artefacts.

#### 2.7.2. Event-related analysis

We epoched the pre-processed data into 3000ms trials (−1000ms to 2000ms relative to stimulus-onset). For each participant, we converted ERPs into percent power change relative to the 500ms pre-stimulus-onset window using the formula: using the formula: (Data – mean baseline values) / mean baseline values.

#### 2.7.3. Time-frequency analysis

Time-frequency representations (TFRs) were computed over 8000ms epochs, (−4000ms to 4000ms relative to stimulus onset). We choose relatively long epochs to avoid edge artefacts due to the transformation. We calculated TFRs by applying a 500ms hanning window as taper on frequencies from 0.5 to 30 Hz in 0.5Hz frequency steps and 5ms temporal steps. Similar to ERPs, we converted participant-specific TFRs into percent power change relative to the 500ms pre-stimulus-onset window.

#### 2.7.4. Intertrial phase coherence estimate (ITPC)

Following time-frequency transformation, we extracted the complex Fourier coefficient for each channel, frequency, and time point in every single trial. Then we computed the phase angles in each trial before finally averaging the single-trial ITPC values over all trials per subject. We performed all analyses in Fieldtrip (Oostenveld et al., 2011; http://fieldtriptoolbox.org).

### 2.8. Statistical Analyses

For all of our analyses, we randomly selected an equal number of events/epochs per condition from N2 and N3 sleep. Evoked events were defined as events that are detected by our algorithms in the 2000ms post-stimulus-onset window. Due to violations to the assumptions of parametric testing, we applied rank-based non-parametric tests using the *nparLD* function implemented in the *nparLD* package available in R (Ngouchi et al., 2012). We report ANOVA-type statistics (ATS), p-values (alpha = 0.05, two-sided), as well as effect sizes using relative treatment effects (RTE). Generally, RTEs represent the probability of the values from the whole dataset being smaller than a randomly chosen observation from the respective group. Therefore, RTE values range between zero and one. An RTE value of 0.5 means no effect. The higher the RTE value of one condition the higher the probability that a randomly chosen value from that condition is larger than that randomly drawn from the whole dataset, and vice versa. When applicable, we performed post-hoc tests via the ‘*NparLD*’ function with Bonferroni’s correction for multiple comparisons. For repeated measures at different time points, we performed non-linear mixed regression via generalized linear mixed models (GLMMs) implemented in the *‘glmer’* function of the *‘lme4’* package in R (Bates et al., 2015). Both KC and micro-arousals were non-normally distributed. Therefore, for K-complexes, we used a GLMM with a Poisson distribution. For micro-arousals, due to the presence of a significant amount of zero counts, we used the zero-inflated Poisson distribution implemented in the ‘*pscl*’ package (Zeileis et al., 2008). We added our subjects as effect with random slopes. We report the estimates of the fixed effects 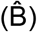 and their standard errors, z-values, and p-values. We performed post-hoc interaction tests using marginal means estimates as implemented in the ‘*emmeans*’ package in R with Tukey’s correction for multiple comparisons and we report Cohen’s d effect sizes.

For a more temporally resolved analysis to compare between the latencies of the detected events, we binned the 2000ms post-stimulus intervals into bins of 100ms, then we calculated the mean of the number of events in each bin for each subject and condition, before finally submitting these results to the permutation analysis in Fieldtrip. The choice of the bin-size is a compromise between a meaningful temporal resolution and a sufficient statistical power.

For ERP, TFR, and ITPC analyses, we selected equal numbers of epochs per condition (66.47 ± 56.25) for each subject. We calculated the grand average over all subjects in each condition before submitting the results to the non-parametric cluster-based permutation analysis in fieldtrip (Maris and Oostenveld, 2007). For this analysis, we selected six frontal channels (F3, F4, F7, F8, Fcz and Fz) and a time window from −500ms to 2000ms relative to stimulus onset. We performed two-sided paired-sample t-tests followed by Monte Carlo’s approximation with 5000 permutations (cluster-alpha = 0.05 and critical alpha = 0.025). We report the sum of the t-values (∑t), as well as Cohen’s d effect sizes calculated over all possible permutations, channels, time points, and frequencies in the cluster.

## 3. Results

### 3.1. Auditory stimuli caused changes to sleep microstructure but not macrostructure

First, we assessed the effects of auditory stimulation on sleep macrostructure. We found that auditory stimulation during sleep does not influence sleep macrostructure. That is, during the experimental night, we found no difference in sleep macrostructure between periods of stimulation and periods of no-stimulation (Table 1). Similarly, there was no change in sleep macrostructure from the adaptation to the experimental night (Table 1-1).

**Table 1.** Sleep macrostructure during stimulation and no-stimulation periods.

|  | TiB (min) | Stimulation (%) | No stimulation (%) | ATS [p] | RTE<sub>stim</sub> |
| --- | --- | --- | --- | --- | --- |
| Wake | 75 [68.87 144.07] | 6.10 [4.56 11.27] | 4.17 [2.74 10.23] | 2.29 [0.13] | 0.55 |
| N1 | 63 [52.33 78.73] | 8.04 [6.59 10.44] | 6.39 [4.41 8.16] | 3.03 [0.08] | 0.58 |
| N2 | 328 [292.89 365.45] | 35.71 [31.3 42.7] | 37.88 [29.5 39.5] | 0.006 [0.94] | 0.50 |
| N3 | 278 [245.34 325.02] | 28.95 [24.8 37.3] | 26.05 [20.6 37.7] | 0.47 [0.49] | 0.54 |
| REM | 161 [139.02 192.63] | 14.39 [12.2 18.8] | 19.92 [14.1 33.1] | 1.001 [0.32] | 0.45 |

\* See also Table 1-1 for a more elaborate description of sleep macrostructure. Values reported medians with 95% confidence intervals. We also report ANOVA-type statistics, *p*-values *npard* package. As effect sizes, we report RTEs of the stimulation condition, the RTE of the no-stimulation can be calculated as follows: $RTE_{nostim} = 1 - RTE_{stim}$ . TiB: Time in bed.

To investigate the effect of the auditory stimulation on sleep microstructure, we compared the numbers of KCs, spindles, and micro-arousals between Stimulus-ON and Stimulus-OFF intervals during the experimental night. Stimulus-ON intervals are 2000ms periods (0 to 2000ms relative to stimulus-onset), while stimulus-OFF intervals are 2000ms windows that start at least 2000ms after the onset of the previous stimulus and during which no sounds are presented (Fig. 4). We found a significantly higher numbers of KCs (*ATS*(*1*) = 54.77, p <0.001, RTE_ON_ = 0.68, RTE_OFF_ = 0.32), slow spindles (*ATS*(*1*) = 9.13, p = 0.002, RTE_ON_ = 0.57, *RTE_OFF_* = 0.43), fast spindles (*ATS*(*1*) = 20.16, p < 0.001, *RTE_ON_* = 0.55, *RTE_OFF_* = 0.45), and micro-arousals (*ATS*(*1*) = 8.45, *p* = 0.003, *RTE_ON_* = 0.58, *RTE_OFF_* = 0.42), in the stimulus-ON than in the stimulus-OFF periods. Moreover, Fig. 4F-H shows the density comparisons, i.e. the number of events per minute of N2 and N3 sleep, between stimulation and no-stimulation periods for KCs, spindles, and micro-arousals.

**Figure 4.**
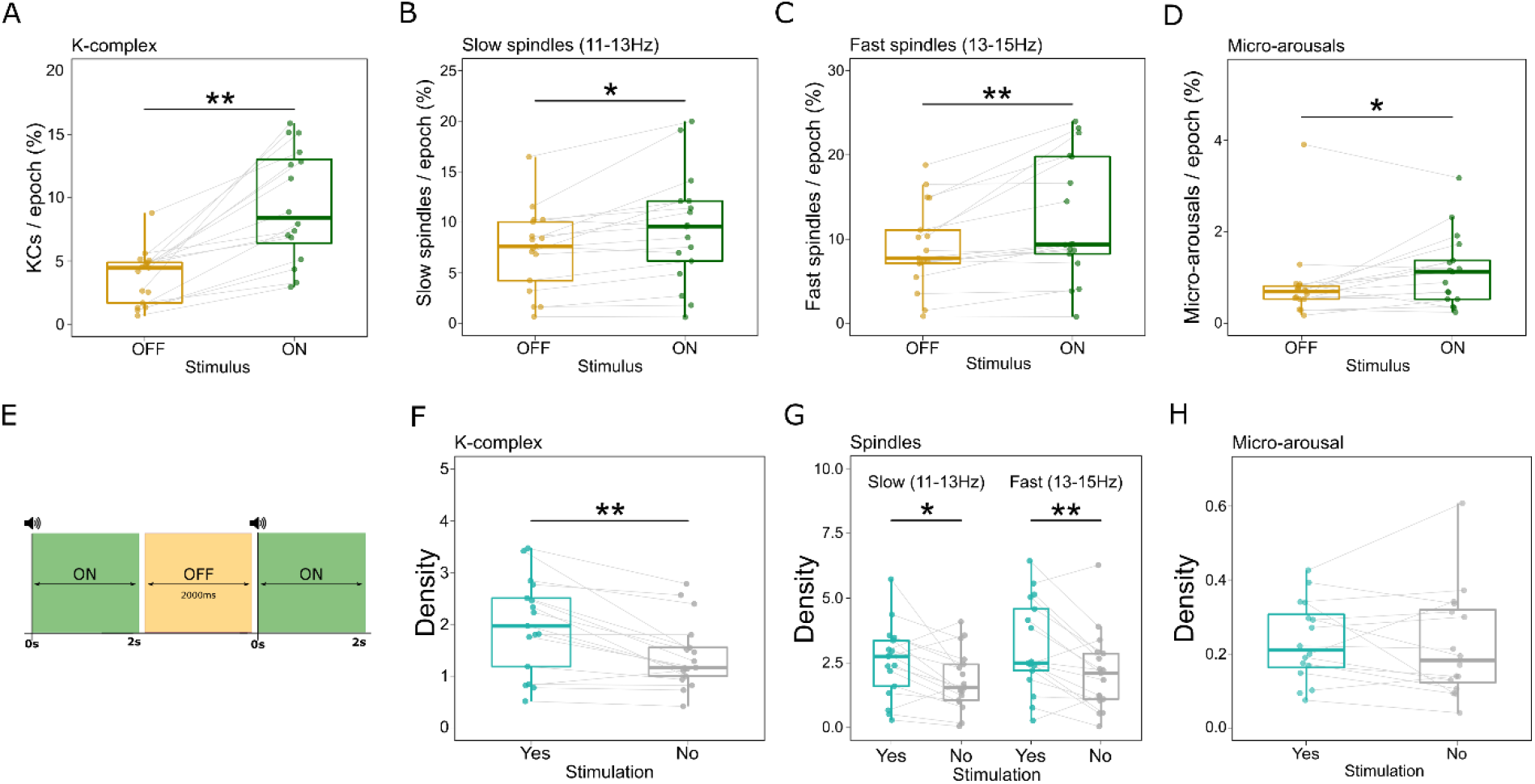
Global effects of auditory stimulation on sleep microstructure. We detected events in the 2000ms post-stimulus window (Stimulus-ON; green), as well as during 2000ms pre-stimulus windows that are at least 2000ms after the last stimulus (Stimulus-OFF, orange). The numbers of A) K-complexes, B) Slow spindles, C) Fast spindles, as well as D) Micro-arousals were significantly higher during the stimulus-ON than the stimulus-OFF periods. Y-axis shows the percentage of detected events in all trials. E) A graphical illustration of the stimulus-ON and stimulus-OFF periods. F-H) Comparison of the density of sleep microstructures during N2 and N3 sleep between stimulation and no-stimulation periods. F) K-complex density during the stimulation periods is higher than that in the no-stimulation periods (ATS(1) = 19.68, p < 0.001, RTE_stim_ = 0.59, RTE_nostim_ = 0.41). G) Slow (ATS(1) = 19.68, p < 0.001, RTE_stim_ = 0.58, RTE_nostim_ = 0.42) and fast (ATS(1) = 8, p = 0.005, RTE_stim_ = 0.58, RTE_nostim_ = 0.42) spindles densities are higher during the stimulation periods. H) micro-arousal density however, did not differ between stimulation and no-stimulation periods (ATS(1) = 1.01, p = 0.31, RTE_stim_ = 0.53, RTE_nostim_ = 0.47). Boxplots show the median and the whiskers depict the 25% and the 75% quartiles. Each dot/triangle represents one participant in one condition. *p<0.05, **p<0.001.

### 3.2. The brain responds selectively to unfamiliar voices during NREM sleep

We subsequently sought to investigate whether brain responses differ depending on the Name and/or Voice used in the stimulus. We employed a non-parametric test with two within factors, i.e. *Name* (SON and UNs) and *Voice* (FV and UFV) from the *nparLD* package. KC responses to auditory stimuli showed a significant effect of voice, as UFVs triggered more KCs than FVs (Fig. 5A, *ATS*(*1*) = 16.10, *p* > 0.001, *RTE_UFV_* = 0.76, *RTE_FV_* = 0.24), no effect of name (*ATS*(*1*) = 0.09, *p* = 0.76, *RTE_SON_* = 0.48, *RTE_UNs_* = 0.52), and a significant interaction *Names x Voices* (*ATS*(*1*) = 11.86, *p* = 0.001). Post-hoc tests revealed that the amount of KCs triggered by the combination FV-SON was marginally higher than that triggered by the combination FV-UNs (*ATS*(*1*) = 4.17, *p_bonf_* = 0.08, *RTE_FVSON_* = 0.56, *RTE_FVUNs_* = 0.44), while there was no difference between UFV-SON and UFV-UNs (*ATS*(*1*) = 4.17, *p_bonf_* = 0.19, *RTE_UFVSON_* = 0.53, *RTE_UFVUNs_* = 0.47).

**Figure 5.**
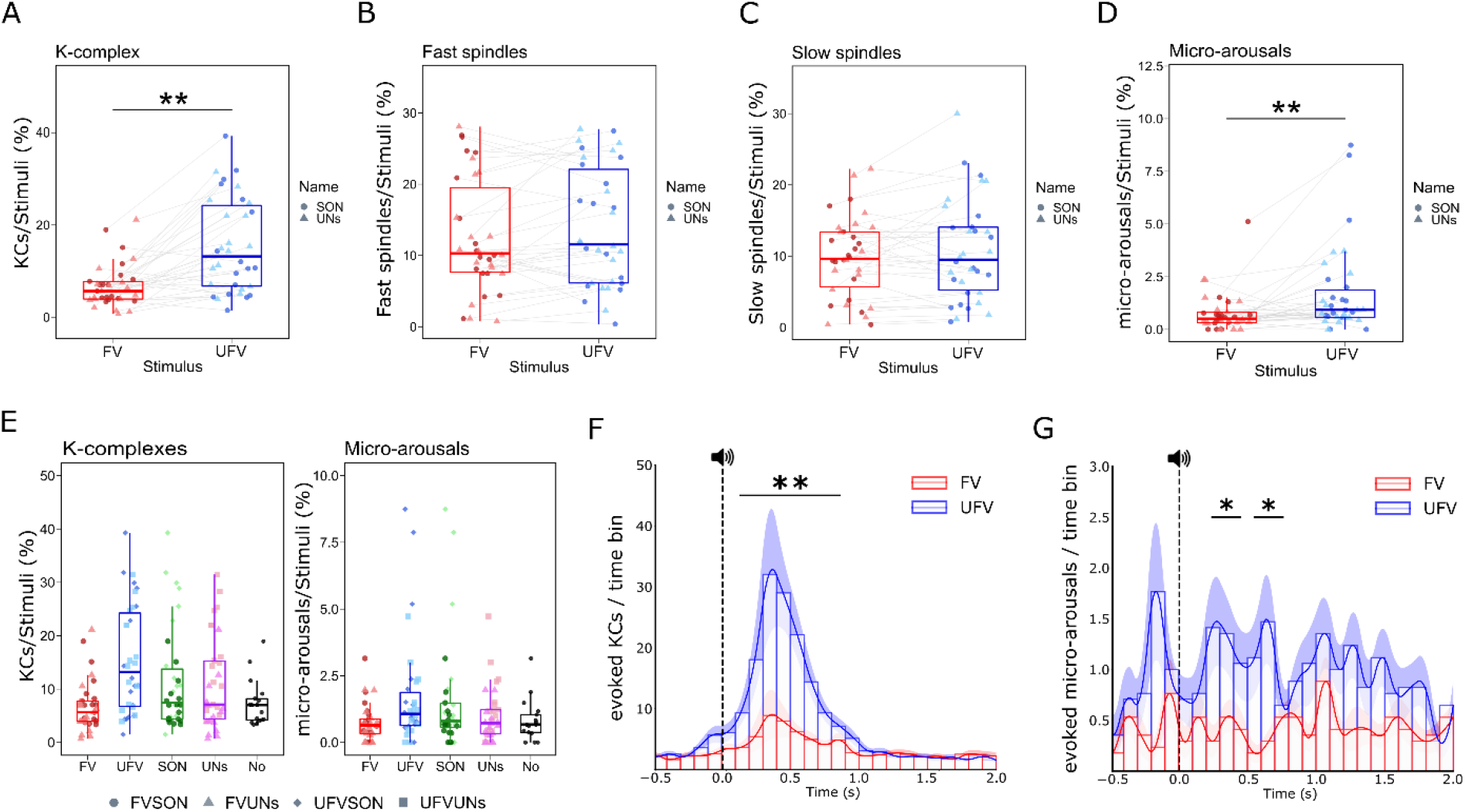
Selective sleep-specific responses to unfamiliar voices during NREM sleep. A) Differences in the triggered K-complexes between familiar voices (FV) and unfamiliar voices (UFV) in the 2000ms post-stimulus-onset window. UFVs triggered more K-complexes than FVs. However, slow (B), and Fast spindles (C) did not differ between FV and UFVs. D) Differences in triggered micro-arousals between FVs and UFVs demonstrating the higher number of micro-arousals that were triggered by UFVs. E) The amount of triggered K-complexes and micro-arousals by all of our stimulus categories as compared to 2000ms no-stimulation epochs. F-G) Temporal aspects of the difference in the triggered K-complexes and micro-arousals, respectively. F) The difference between UFV and FV in the number of triggered K-complexes was significant from 100ms to 800ms (indicated by bar and asterisks). G) The difference in micro-arousals between FVs and UFVs. We found a significant difference in the number of triggered micro-arousals in the periods from 200 to 400ms, and from 500 to 700ms. Boxplots show the median and the whiskers depict the 25% and the 75% quartiles. Each dot/triangle represents one participant in one condition. The lines depicts the means in each bin and the shadings show the standard error of the mean. *p<0.05, **p<0.001, SONs: Subject’s own name, UNs: Unfamiliar names. FVSON: familiar voice speaking the subject’s own name. UFVSON: unfamiliar voice speaking the subject’s own name, FVUNs: familiar voice speaking two unfamiliar names, and UFVUNs: unfamiliar voice speaking two unfamiliar names.

For fast and slow spindles, there was no effect of voice (Fig. 5B-C, fast: *ATS*(*1,16*) = 2.71, *p* = 0.10, *RTE_UFV_* = 0.53, *RTE_FV_* = 0.47 - slow: *ATS*(*1,16*) < 0.001, *p* = 0.99, *RTE_UFV_* = 0.5, *RTE_FV_* = 0.5), or name (fast: *ATS*(*1,16*) = 0.62, *p* = 0.43, *RTE_SON_* = 0.49, *RTE_UNs_* = 0.51 - slow: *ATS*(*1,16*) = 0.46, *p* = 0.5, *RTE_SON_* = 0.51, *RTE_UNs_* = 0.49), and no interaction Names X Voices (fast: *ATS*(*1,16*) = 2.63, *p* = 0.10, *RTE_UFVSON_* = 0.50, *RTE_UFVUNs_* = 0.55, *RTE_FVSON_* = 0.47, *RTE_FVUNs_* = 0.47, slow: *ATS*(*1,16*) = 0.11, *p* = 0.74, *RTE_UFVSON_* = 0.51, *RTE_UFVUNs_* = 0.49, *RTE_FVSON_* = 0.52, *RTE_FVUNs_* = 0.48).

Micro-arousals showed a main effect of voice (Fig. 5D, *ATS*(*1*) = 9.14, *p* = 0.002, *RTE_UFV_ = 0.59, RTE_FV_* = 0.41), no effect of name (*ATS*(*1*) = 1.16, *p* = 0.29, *RTE_SON_* = 0.53, *RTE_UNs_* = 0.47) and no interaction Names x Voices (*ATS*(*1*) = 0.19, *p* = 0.66, *RTE_UFVSON_* = 0.61, *RTE_UFVUNs_* = 0.58, *RTE_FVSON_* = 0.44, *RTE_FVUNs_* = 0.37).

Next, we performed a more temporally-resolved analysis (see section 2.8), and found that the difference in the evoked KCs between FVs and UFVs occurred as early as 100ms post-stimulus-onset and lasted until 800ms post-stimulus-onset (Fig. 5F, ∑*t*(*16*) = 25.35, *p* = 0.002, *d* = 1.02). For micro-arousals (Fig. 5G), the difference appeared 200ms post-stimulus-onset, and remained significant between 200ms and 700ms post-stimulus (200-400ms: *t*(*16*) = 5.92, *p* = 0.01, *d* = 0.84, 500-700ms: *t*(*16*) = 5.19, *p* = 0.02, *d* = 0.76).

To ask whether the unfamiliarity of the voice principally accounts for the difference in KC and micro-arousal responses between voices, we divided the night into halves and hypothesized that brain responses to UFVs, but not to FVs, will decrease from the first to the second half (Fig. 6). To illustrate this, we modelled the change in KCs using a GLMM with Poisson distribution fit by maximum likelihood (Laplace Approximation) (Fig. 6A). We found an effect of Voice (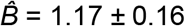, *z* = 7.51, *p* < 0.001), no effect of Time (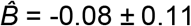, *z* = −0.77, *p* = 0.44), and a significant interaction Times X Voice (z = −2.28, *p* = 0.02). Post-hoc tests revealed a significant decrease for UFV-triggered KCs from the first to the second half of the night (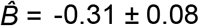, *z* = 3.54, *p* = 0.002, *d* = 0.31), while FV-evoked KCs did not change (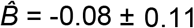, *z* = 0.77, *p* = 0.87, *d* = 0.08). For micro-arousals, a GLMM with zero-inflated Poisson distribution (Fig. 6B) demonstrated a marginally significant effect of Voice (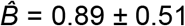, *z* = 1.75, *p* = 0.08), no effect of Time (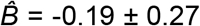, *z* = 0.73, *p* = 0.98), and no interaction Times X Voice (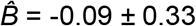, *z* = −0.27, *p* = 0.55). We provide a more detailed description of the GLMMs results in Fig. 6C-D.

**Figure 6.**
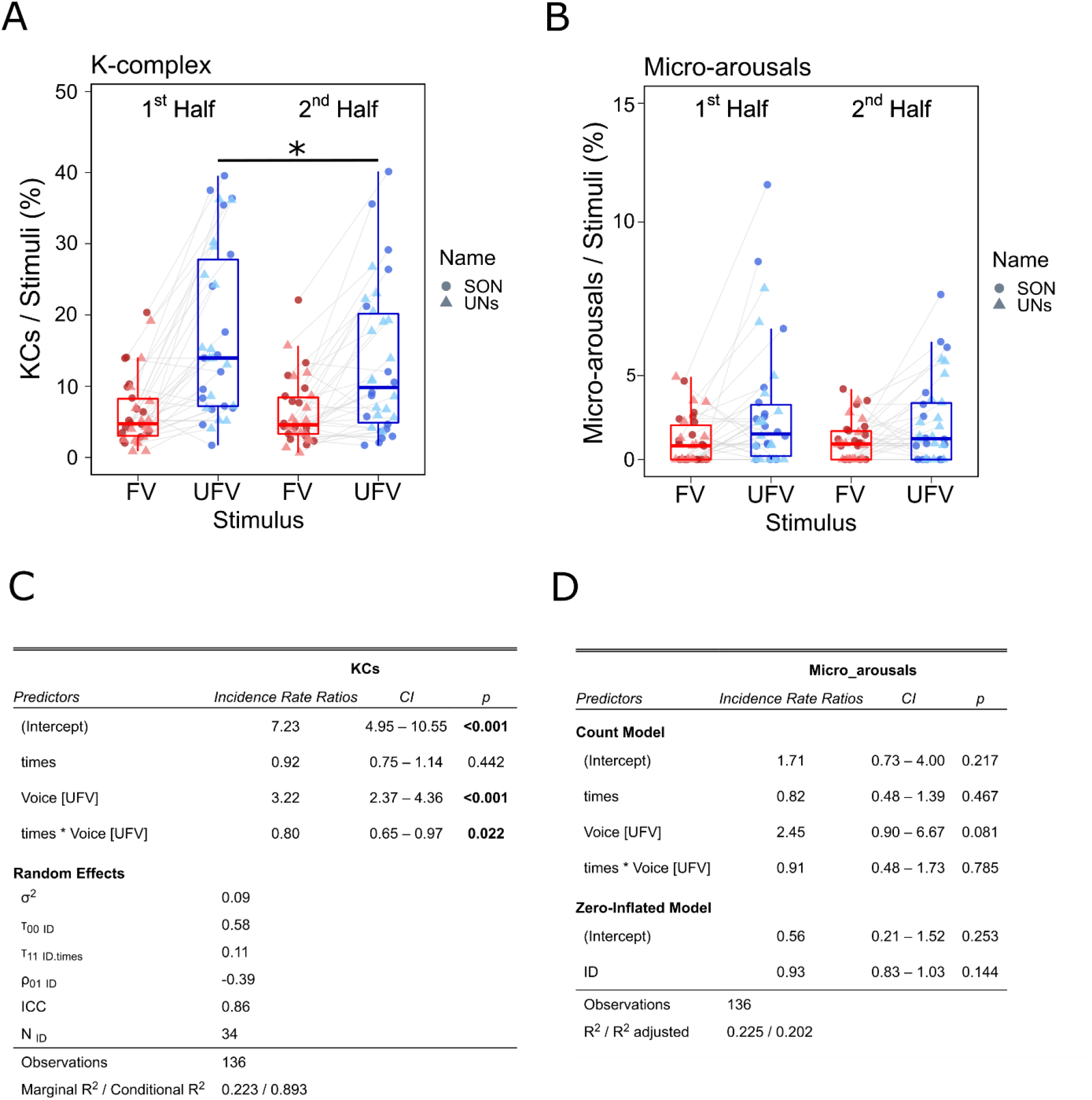
The effect of time on sleep-specific responses to unfamiliar voices during NREM sleep. A) The difference in the numbers of triggered K-complexes to familiar voices (FV) and unfamiliar voices (UFV). A GLMM using Poisson distribution revealed a significant interaction Time x Voice, as only the amount of UFV-triggered K-complexes decreased in the second half. B) The number of triggered micro-arousals during the first and the second half of the night. C) Statistical details of the GLM model of K-complexes. We added Time (first half & second half), and Voices (familiar and unfamiliar voices) as fixed effects. Moreover, we assigned a random slope for each subject. We found a significant main effect of voice, no effect of time, and a significant interaction Time X Voice. D) Statistical details of the zero-inflated Poisson GLM of micro-arousals. Similar to K-complexes, we added Time (first half & second half), and Voices (familiar and unfamiliar voices) as fixed effects and random slopes for each subject. No main effect of time and no interaction Time x Voice, indicating that the amount of triggered micro-arousals did not change from the first to the second half of the night. *p<0.05, SONs: Subject’s own name, UNs: Unfamiliar names. Boxplots show the median and the whiskers depict the 25% and the 75% quartiles. Each dot/triangle represents one participant in one condition.

### 3.3. UFVs evoke stronger and better time-locked brain responses during NREM sleep

Next, we aimed to examine the neural dynamics underlying the aforementioned differences in sleep microstructure. In a first analysis step, we compared the ERPs between FVs and UFVs in two conditions: 1) when the stimuli trigger KCs, 2) when no KCs are triggered. We found that when KCs are triggered, UFVs evoked a larger, more pronounced negative wave that resembles the KC’s N550 component (Fig. 7A, ∑*t*(*16*) = −436.35, *p* > 0.001, *d* = 0.96). We thought of two possible explanations for this difference in the ERPs in the first condition. One explanation might be that UFV-evoked KCs have larger amplitudes relative to FV-evoked KCs. Another possibility would be a stronger time locking of the evoked responses to UFVs as compared to FVs. To disentangle this, we compared the peak-to-peak amplitudes of the KCs detected by our algorithm, and investigated the phase consistency across trials trial types (FV vs. UFVs) via inter-trial phase coherence (ITPC, Tallon-Baudry et al. 1996). We found no difference in the amplitudes of the evoked KCs between voices (Fig. 7B, *ATS*(*1,16*) = 1.47, *p* = 0.23, *RTE_UFV_* = 0.48, *RTE_FV_* = 0.52), or names (*ATS*(*1,16*) = 0.71, *p* = 0.39, *RTE_SON_* = 0.52, *RTE_UNs_* = 0.48) as well as no interaction (*ATS*(*1,16*) = 1.47, *p* = 0.23, *RTE_UFVSON_* = 0.49, *RTE_UFVUNs_* = 0.47, *RTE_FVSON_* = 0.54, *RTE_FVUNs_* = 0.50). Fig. 7C shows the ITPC contrast between FVs and UFVs indicating that difference in the IPTC is driven mainly by stronger time-locking to UFVs. However, we observed stronger ITPC for the UFV condition as compared to FV presentations in lower frequencies (1-4Hz) (Fig 7D, ∑*t*(*16*) = 2413.98, *p* = 0.003, *d* = 1.06) indicating a more consistent timing of brain responses evoked by the UFV stimulation. Interestingly, when no KCs are evoked, there was no difference in the amplitude of the ERPs (Fig. 7B, 0.3 - 0.37s: ∑*t*(*16*) = 27.31, *p* = 0.13), nor in IPTC values (Fig. 7F, ∑*t*(*16*) = 2521.7, *p* = 0.001) between FVs and UFVs

**Figure 7.**
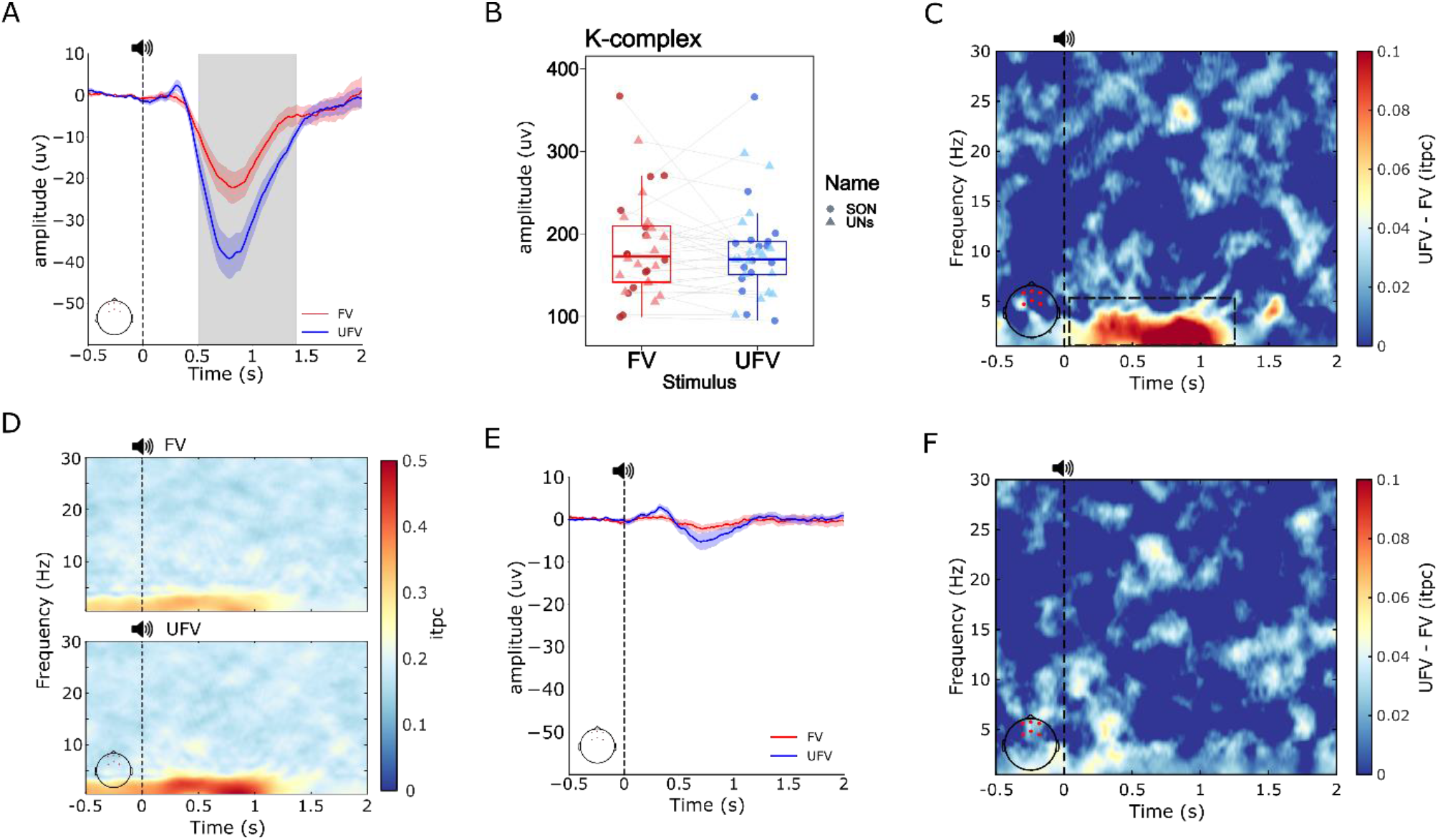
Brain responses in the presence of K-complexes are stronger and better time-locked to unfamiliar voice stimuli. A) ERP contrast between KC-triggering UFV and FV stimuli. UFV triggered a larger amplitude of the evoked response around the same time window of the N550 component of the K-complex. The grey shadings represent the time range that shows a significant difference (510ms to 1400ms). B) Comparison of the peak-to-peak amplitude showing no difference in the amplitudes of the evoked K-complexes between FVs and UFVs. C) The difference in ITPC values between UFVs and FVs in the presence of K-complexes. UFVs evoked significantly higher ITPC than FV in the delta (1-4Hz) frequency band. Note that the largest difference in phase locking overlaps with the difference in the evoked response (~500-1000ms). D) Separate ITPC plots for Familiar voices (FVs, top) and unfamiliar voices (UFVs, bottom). E) ERP responses to FVs and UFVs in the absence of the evoked K-complexes. There was no difference in amplitudes between FVs and UFVs when no K-complexes were evoked. F) ITPC difference between UFVs and FVs in the absence of the evoked K-complexes. There is no difference in ITPC between voices in the absence of K-complexes. Vertical dashed lines (at x= 0) represent stimulus onset. Blue and red lines represent the mean at each time point and the shadings represent the standard error of the mean. The locations of the channels used for the analysis are indicated by the red dots on the topographical plots at the bottom left side of the plots.

### 3.4. Oscillatory responses to different voices during NREM sleep

Finally, we contrasted oscillatory brain responses between UFVs and FVs. We found that UFVs always triggered stronger power in lower frequencies, i.e. delta range (1-4Hz) starting around 250ms post stimulus onset, independent of the presence (Fig. 8A, ∑*t*(*16*) = 9395.64, *p* < 0.001, *d* = 1.25), or absence (Fig. 8B, ∑*t*(*16*) = 4097.56, *p* = 0.013, *d* = 0.86) of the evoked KCs. However, only in the presence of KCs, UFVs additionally elicited a significant increase in oscillatory power at higher frequencies as compared to FVs (Fig. 8A, ∑*t*(*16*) = 5020.62, *p* = 0.006, *d* = 1.04), indicative of a (stronger) arousal response to UFVs relative to FVs.

**Figure 8.**
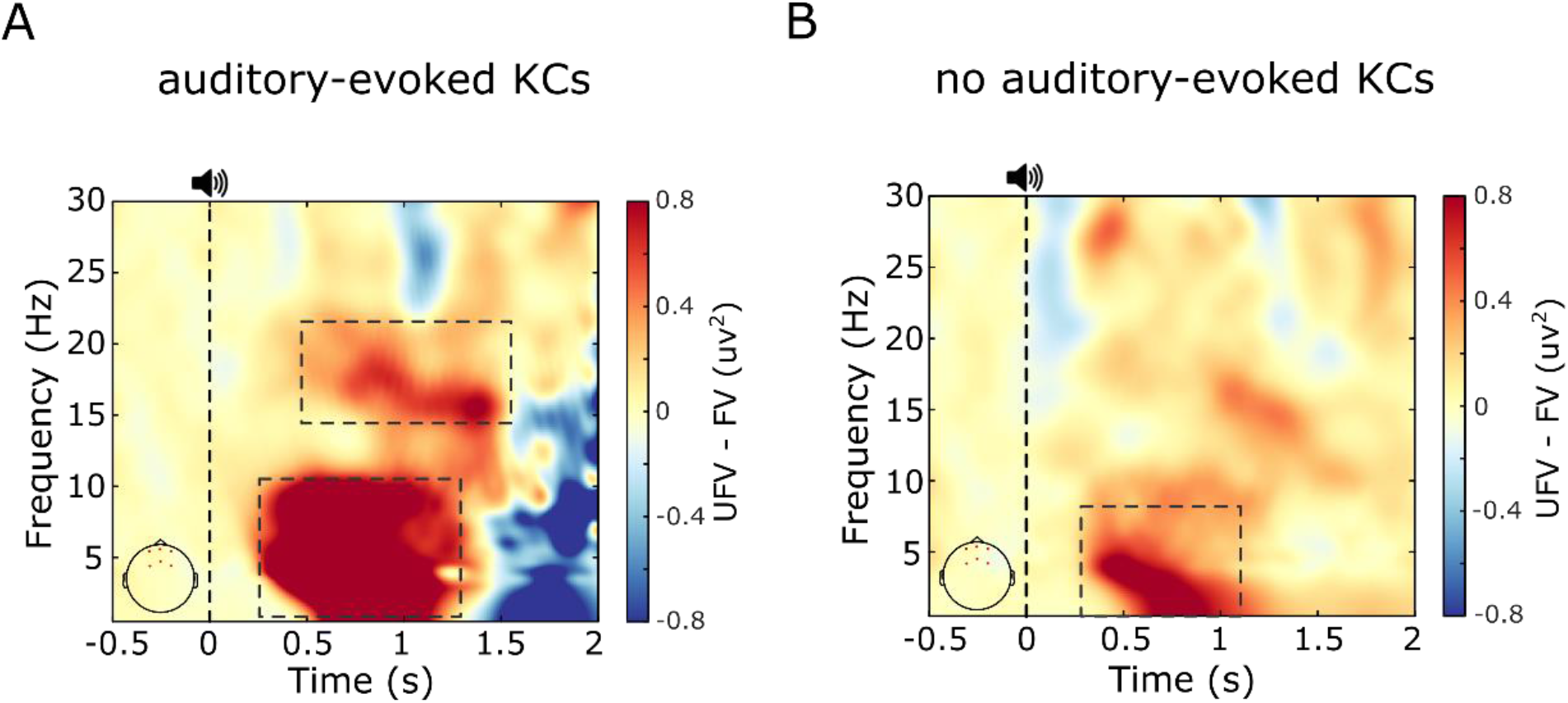
High frequency oscillatory responses are higher to UFVs relative to FVs in the presence of K-complexes. Spectral maps of the oscillatory power differences between familiar voices (FV) and unfamiliar voices (UFV) demonstrate stronger oscillatory responses to UFV in a broad frequency range (~1-10Hz) regardless of the presence (A) or the absence (B) of K-complexes. However, stronger high frequency (>16Hz) responses to UFVs appeared only in the presence of K-complexes. Vertical dashed lines (at x= 0) represent stimulus onset. The locations of the channels used for the analysis are indicated by the red dots on the topographical plots at the bottom left side of the plots.

## 4. Discussion

In this study, we presented sleepers with their own first names (SON) and two unfamiliar names (UNs) spoken by either a familiar voice (FV) or an unfamiliar voice (UFV) during a full night of sleep with polysomnography. We show that although auditory stimuli did not cause any changes in sleep architecture (Table 1), they induced prominent, stimulus-specific changes in sleep microstructure. Generally, presenting auditory stimuli during NREM sleep increased the number of triggered KCs, spindles, and micro-arousals (Fig. 4). The (paralinguistic) stimulus content influenced its ability to elicit KCs and micro-arousals whereas spindles were not affected. UFVs triggered more KCs (Fig. 5A) than FVs with the number of the UFV-triggered K-complexes decreasing in the second half of the night (Fig. 6A). Similarly, UFVs triggered more micro-arousals (Fig. 5D) than FVs. However, we found no difference in the amount of triggered KCs, spindles, or micro-arousals between SONs and UNs. A more fine-grained investigation indicated that in the presence of the auditory-evoked KCs, UFVs triggered larger, more-synchronized brain responses as compared to FVs (Fig. 7A-C). These differences vanished when comparing UFV and FV stimuli that did not evoke KCs (Fig. 7E-F). Similarly, oscillatory responses to UFVs demonstrated stronger high-frequency arousal responses (>16 Hz) only in the presence of the evoked KCs (Fig. 8).

It has previously been suggested that the more relevant the stimulus, the higher its tendency to trigger KCs (Halász, 2005). In this regard, our results pose UFVs as more relevant - or in evolutionary terms potentially more threatening (Blume et al., 2018) – and consequently more arousing to the sleeper than FVs. Indeed, the increase in micro-arousals following UFVs suggests a transient shift towards external processing of “vital” environmental stimuli. Crucially, the notion that UFVs, but not FVs, triggered less KCs in the second half than in the first half of the night (Fig. 6A) cannot be fully explained by stimulus-specific adaptation processes, as the number of KCs elicited by FVs remained unchanged. Rather, such a finding might support the notion that the sleeping brain continues to learn new information during sleep (Züst et al., 2019). That is, the brain learns that an initially unfamiliar stimulus posed no immediate threat to the sleeper and therefore decreases its response to it. Conversely, in a normal and safe sleep environment, the sleeping brain might be “expecting” to hear FVs and consequently attenuates its responses to such stimuli. Although this notion remains speculative, it entails a qualitative investigation of the brain’s ability to generate predictions of the external sensory environment during sleep. Taken together, our results suggest that the unfamiliarity of a voice is a strong promoter of brain responses during NREM sleep.

Another crucial point that we attempted to clarify in this study is the role of the auditory-evoked brain responses during NREM sleep. Central to such responses is the KC, i.e. the most prominent sleep-specific response to sensory stimulation. To do so, we contrasted trials during which FVs and UFVs triggered KCs. In such trials, UFVs evoked a larger negative component that resembles the N550 component of the KC (Fig. 7A). The N550 has been associated with large-scale neuronal silencing that protects sleep (Cash et al., 2009; Laurino et al., 2014), and conversely, an arousal reaction that facilitates stimulus processing (Atienza et al., 2001). In our study, the difference in the N550-amplitude between FVs and UFVs overlapped temporally with the stronger ITPC to UFVs (500 - 1000ms post-stimulus; Fig. 7C). The stronger ITPC to UFVs reflects better temporal synchronization of brain responses to UFVs and therefore supports the notion of a transient window for information processing. That is, stimulus-induced phase modulations have been suggested to promote information processing and transmission in the cortex (Canavier, 2015; Lakatos et al., 2013; Voloh and Womelsdorf, 2016) and increased ITPC values have been associated with better cognitive performance (Hanslmayr et al., 2005; Eidelman-Rothman et al., 2019) and enhanced attention (Joon Kim et al., 2007) during wakefulness. Importantly, the narrower peak for the evoked KCs following UFVs (Fig. 5F), which reflects the grouping of the evoked KCs within a shorter temporal window, further suggests a contribution for KCs in the observed phase modulations. In the same vein, our oscillatory analysis corroborates a role for the auditory-evoked KCs in sensory processing. UFVs triggered stronger high frequency (>16Hz) responses relative to FVs, which appeared only in the presence of KCs (Fig. 8), suggesting an arousal-like response. Together, our findings suggest a central role for KCs in the extraction and processing of relevant auditory information during NREM sleep.

It is worth noticing that, in our study, the peak of the negative component occurred later (~ 750-800ms) than the usual time window of the N550, i.e. 500-550ms, and with a much smaller amplitude (20 - 50μv) than that observed (~100μv) in previous literature (Colrain, 2005; Halász, 2005). The increased latency of the N550 might be due to the relatively large temporal window we used to detect evoked KCs (2000ms post-stimulus) as well as the longer mean duration of the stimuli (~808ms). The smaller amplitude, however, we trace back to our choice of an average reference as compared to the mastoids reference used in earlier studies (Bastien and Campbell, 1992; 1994; Laurino, 2014).

Contrary to the results of Perrin et al. (1999), we observed no difference in brain responses between SON and UNs. It is plausible that our methodological approach, for example the choice of an average references, might have mitigated subtle brain responses leading to such discrepancy in the results. Indeed, while sleep preserves low-level auditory processing, it attenuates higher-order linguistic tracking (Makov et al., 2017). We speculate that this is due to the disruption of the activity of the large-scale networks necessary for higher-order name processing (Démonet et al., 1992; Luke et al., 2002), which might be due to the loss of long-range cortical connectivity during sleep (Massimini et al., 2005). Further research should aim to elucidate the mechanisms of language tracking during sleep.

We show that the auditory-evoked spindles are not influenced by specific characteristics of the auditory stimuli (i.e., name or voice). However, the role of spindles in response to sensory stimulation during sleep is far from clear. Previous research has shown that spindles attenuate or even inhibit the processing of auditory information (De Gennaro and Ferrera, 2003; Schabus et al., 2012; Blume et al., 2018). Other work suggests that brain responses are preserved during spindles (Sela et al., 2016) and even argues for a role for spindles in the processing of memory-related sensory information (Antony et al. 2018; Cairney et al. 2018; Jegou et al. 2019). More research should investigate the role of spindles in response to sensory information irrelevant to ongoing memory processes.

Finally, we found no relevant changes in sleep macrostructure and architecture due to auditory stimulation (Table 1). In fact, micro-arousals represent an integral part of healthy sleep as they tailor sleep processes according to internal and external demands thus ensuring its reversibility (Halász et al., 2004). Further, the analysis of microstates even indicates a shift towards deeper sleep in response to UVF stimuli (Table1-1C), which is most likely a by-product of more auditory-evoked KCs that influenced the staging. This is in line with previous literature that suggests a dual function of KCs (Halász, 2005, 2016; Jahnke et al., 2012; Blume et al., 2017, 2018; Legendre et al., 2019; Latreille et al., 2020). However, the classical 30s sleep staging might not be sensitive enough to capture subtle changes in sleep microstructure. Therefore, the development and refinement of new fine-grained methods, such as the Hori-based microstate classification (Jagannathan et al., 2018), promises better monitoring of transient sleep fluctuations, especially in the presence of sensory perturbation.

In summary, sleep is far from being a homogenous state of unconsciousness. There exist windows in sleep during which the brain filters, extracts, and even processes relevant information. We speculate that such content-specific, dynamic reactivity to incoming sensory information enables the brain to enter a sentinel processing mode (Blume et al., 2018) in which it preserves its ability to efficiently engage in the many important processes that are ongoing during sleep while remaining connected to the surrounding environment.

## 5. Acknowledgments

This project is supported by the Austrian Science Fund FWF (Y777). MS Ameen is supported by the FWF Austrian Science Fund (W 1233-B) and the Austrian Academy of Science (OEAW). The authors would like to thank Renata del Guidice for her participation in data collection, Kerstin Hoedlmoser for her support, and the members for the Sleep lab Salzburg for their invaluable input throughout the process.

