## Supplementary material for "The brain selectively tunes to unfamiliar voices during sleep": Table 1-1

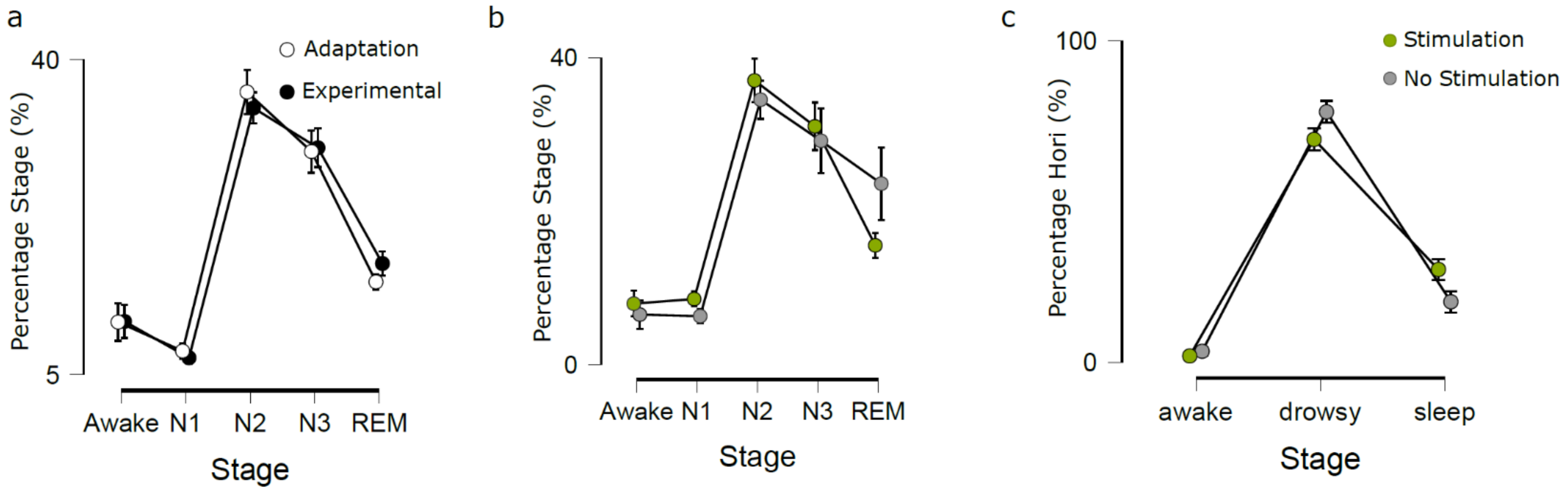


***Table1-1. The influence of auditory stimulation on sleep macrostructure.*** *A) Difference in sleep architecture, i.e. the distribution of sleep stages between the adaptation and the experimental nights. We selected equal number of epochs for both conditions. To account for the sleep-onset-latency effect we selected epochs starting from the first N1 epoch and we discarded all awake epochs before that point. A non-parametric test with two-with factors: night (adaptation vs experimental) and sleep stage (Awake, N1, N2, N3, and REM) revealed an effect of sleep stage (ATS(2.61) = 62.75, p < 0.001, $RTE_{wake}$ = 0.24, $RTE_{N1}$ = 0.26, $RTE_{N2}$ = 0.79, $RTE_{N3}$ = 0.70, $RTE_{REM}$ = 0.51). There was no effect of the night (ATS(1) = 0.04, p = 0.84, $RTE_{Nostim}$ = 0.48, $RTE_{stim}$ = 0.52) and no interaction night X stage (ATS(2.56) = 0.75, p = 0.51). B) Difference in sleep architecture between the stimulation and no stimulation periods within the experimental night. We selected equal number of epochs for both conditions starting from the first N1 epoch. We found a main effects for stage (ATS(2.51) = 53.30, p < 0.001, $RTE_{wake}$ = 0.28, $RTE_{N1}$ = 0.18, $RTE_{N2}$ = 0.82, $RTE_{N3}$ = 0.74, $RTE_{REM}$ = 0.47). There was no effect of the stimulation (ATS(1) = 3.36, p = 0.07, $RTE_{ADAPT}$ = 0.5, $RTE_{EXP}$ = 0.5), and no interaction stimulation X stage (ATS(2.08) = 0.68, p = 0.52). C) Microstate differences between the stimulation and the no stimulation conditions. To circumvent the poor temporal resolution of classical sleep staging in 30sec epochs, we opted for a more time-resolved (four-second epochs) analysis of sleep stages based on the Hori scoring system. We found a main effects for stage (ATS(1.64) = 146.26, p < 0.001, $RTE_{wake}$ = 0.28, $RTE_{N1}$ = 0.18, $RTE_{N2}$ = 0.82, $RTE_{N3}$ = 0.74, $RTE_{REM}$ = 0.47). There was no effect of the stimulation (ATS(1.69) = 3.36, p = 0.19, $RTE_{ADAPT}$ = 0.5, $RTE_{EXP}$ = 0.5), and a significant interaction stimulation X stage (ATS(1.47) = 8.17, p = 0.02). Post-hoc pairwise test with Bonferroni's correction for multiple comparisons revealed that auditory stimulation resulted in higher number of sleep epochs (ATS(1) = 12.84, p < 0.001, $RTE_{stim}$ = 0.62, $RTE_{nostim}$ = 0.38) and lower number of drowsy epochs (ATS(1) = 10.88, p < 0.001, $RTE_{stim}$ = 0.4, $RTE_{nostim}$ = 0.6), suggesting deeper sleep during stimulation periods which simply might be a by-product of the auditory-evoked k-complexes. Dots represent the means over subjects and the error bars represent the standard error of the mean.*
